# Nucleosides drive histiocytosis in SLC29A3 disorders by activating TLR7

**DOI:** 10.1101/2019.12.16.877357

**Authors:** Takuma Shibata, Masato Taoka, Shin-Ichiroh Saitoh, Yoshio Yamauchi, Yuji Motoi, Mayumi Komine, Etsuko Fujita, Ryota Sato, Hiroshi Sagara, Takeshi Ichinohe, Mimi Kawazoe, Chiharu Kato, Katsuhiro Furusho, Yusuke Murakami, Ryutaro Fukui, Mamitaro Ohtsuki, Umeharu Ohto, Toshiyuki Shimizu, Nobuaki Yoshida, Toshiaki Isobe, Kensuke Miyake

## Abstract

A lysosomal transmembrane protein SLC29A3 transports nucleosides from lysosomes to the cytoplasm. Loss-of-function mutations of the SLC29A3 gene cause lysosomal nucleoside storage in monocyte/macrophages, leading to their accumulation called histiocytosis in humans and mice. Little is known, however, about a mechanism behind nucleoside-dependent histiocytosis. TLR7, an innate immune sensors for single stranded RNA, bind and respond to nucleosides. We here show that they drive nucleoside-mediated histiocytosis. Patrolling monocyte/macrophages accumulate in the spleen of *Slc29a3*^−/−^ mice but not *Slc29a3*^−/−^ *Tlr7*^−/−^ mice. Accumulated patrolling monocyte/macrophages stored nucleosides derived from cell corpse. TLR7 was recruited to phagosomes and activated as evidenced by TLR7-dependent phagosomal maturation. TLR7 induced hyper-responsiveness to M-CSF in *Slc29a3*^−/−^ monocyte/macrophages. These results suggest that TLR7 drives histiocytosis in SLC29A3 disorders.

**One Sentence Summary:** SLC29A3 disorders are caused by activation of TLR7 with accumulated nucleosides in lysosomes.

## Main Text

Our recent results revealed an unexpected link between Toll-like receptors (TLRs) and some forms of histiocytosis, a disease characterized by expansion of dendritic cells or monocyte/macrophages (*1, 2*). TLRs serve as pathogen sensors in monocytes and macrophages. TLR7 and 8 respond to pathogen-derived RNA (*3, 4*) and their excessive activation drives macrophage activation syndrome in mice (*5–8*), although underlying mechanisms are not completely understood. The structures of TLR7 and 8 indicate that nucleosides act on TLR7 and 8 (*9–11*). TLR7 binds to guanosine (Guo) or 2’-deoxyguanosine (dGuo) as well as to uridine (Uri)-containing ORN, whereas TLR8 interacts with Uri and purine-containing ORN. Very interestingly, abnormal nucleoside storage is known to cause some monogenic forms of histiocytosis, although molecular mechanisms behind were not understood. Our recent findings on the interaction between TLR7/8 and nucleosides could be applicable to such human diseases.

It was previously known that RNA degradation in endosome/lysosomes proceeds up to nucleosides, which are transported to the cytoplasm for further degradation into the end-metabolite uric acid. SLC29A3, a lysosomal membrane protein abundantly expressed in monocyte/macrophages, mediates nucleoside transport across the lysosomal membranes (*12*). Mutations of the SLC29A3 gene cause monogenic diseases including H syndrome, Faisalabad Histiocytosis, pigmented hypertrichosis with insulin-dependent diabetes mellitus (PHID) syndrome, and familial Rosai-Dorfman disease (*13–15*). These SLC29A3 disorders are characterized by histiocytosis due to nucleoside storage in endosome/lysosomes (*13–15*). However, mechanisms by which nucleoside storage causes histiocytosis have not been understood. We hypothesized that nucleosides such as Guo act on TLR7 in SLC29A3 disorders to drive histiocytosis.

*Slc29a3*^−/−^ and *Tlr7*^−/−^ mice were generated and crossed with each other to generate *Slc29a3*^−/−^*Tlr7*^−/−^ mice (Fig. S1, S2). The organs of *Slc29a3*^−/−^ mice were examined by Liquid Chromatography-Mass Spectrometry (LC-MS) to evaluate nucleoside accumulation. While tissue levels of Guo and dGuo were difficult to detect in wild-type mice, we found them in the spleen of lysosomal nucleoside transporter-deficient *Slc29a3*^−/−^ mice (Fig. 1A, Fig. S3). As human and mouse TLR7s respond to Guo and dGuo, but not other nucleosides (9), Guo and dGuo that accumulated in *Slc29a3*^−/−^ mice may act on TLR7. Consistent with the previous report (*16*), spleens of *Slc29a3*^−/−^ mice were large due to increases in cell number (Fig. 1B, Fig. S4A). Concerning cell type-specific changes in spleen and peripheral blood, increases in the percentages were restricted to CD11b^+^Ly6G^−^ monocyte/macrophages (Fig. 1C-D. Fig. S4B-G) and especially a subset thought to patrol the vasculature and clear damaged endothelial cells in the circulation (Fig. 1E, F, S4H) (*17*). We hereafter describe CD11b^+^ Ly6C^lo^ FcγRIV^hi^ cells accumulated in *Slc29a3*^−/−^ mice as patrolling monocytes/macrophages. These changes were totally dependent on TLR7, as demonstrated in *Slc29a3*^−/−^ *Tlr7*^−/−^ mice (Fig. 1B-F, S4). These results indicate that TLR7 drives histiocytosis in *Slc29a3*^−/−^ mice.

**Fig. 1.**
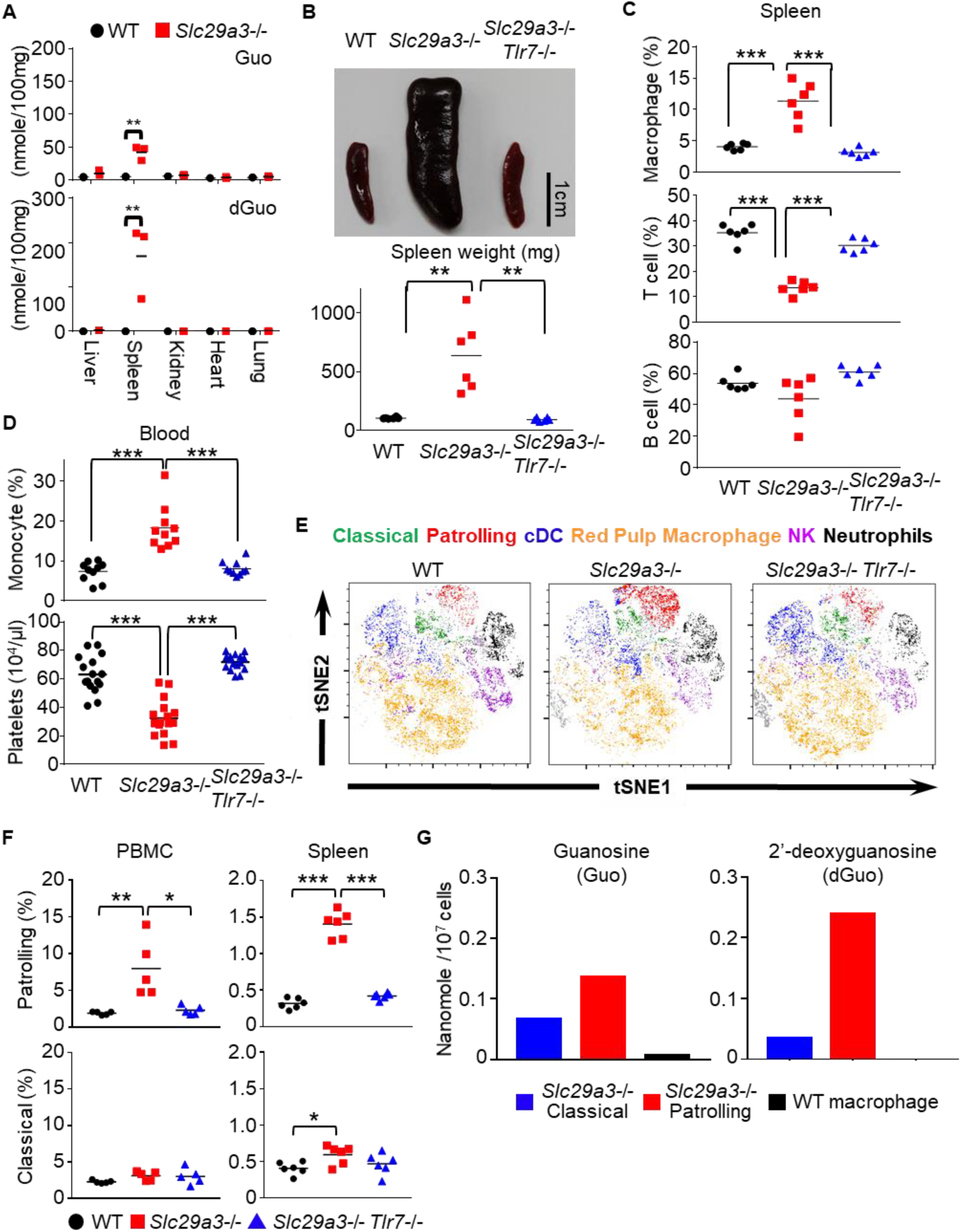
TLR7-dependent histiocytosis in *Slc29a3*^−/−^ mice. **A**, The amount of Guanosine (Guo) and 2’-deoxyguanosine (dGuo) in various organs (nanomole/100 mg tissue). Each dot shows the value obtained from one mouse. n=3 biological replicates (n=3). **B**, Representative gross appearance of spleens from 6-month-old mice (top). scale bar, 1 cm. The bottom graph shows spleen weight from wild-type (WT), *Slc29a3*^−/−^ and *Slc29a3*^−/−^ *Tlr7*^−/−^ mice (n=6). **C**, The percentage of splenic CD11b^+^Ly6G^−^ monocytes/macrophages, CD19^+^ B cells, and CD3ε^+^ T cells from 6 months old mice was determined by flowcytometric analysis (n=6). **D**, The percentage of CD11b^+^ Ly6G^−^ monocytes (n=10) and the platelet counts (n=16) in peripheral blood from 2∼3 months old mice. **E**, t-distributed stochastic neighbor embedding (t-SNE) plot of CD11b^+^ splenocytes analyzed by flowcytometry. Colored cell types were determined by the differential expression level of known markers. Classical macrophages (Ly6G^−^ NK1.1^−^ Ly6C^+^ FcγRIV^hi^ IA/IE^low^), Patrolling macrophages (Ly6G^−^ NK1.1^−^ Ly6C^−^ FcγRIV^hi^ IA/IE^low^), cDC (Ly6G^−^ NK1.1^−^ CD11c^+^ Ly6C^−^ IA/IE^high^), Red pulp macrophages (F4/80^+^ High autofluorescence), NK cells (NK1.1^+^), and Neutrophils (Ly6G^+^) **F**, The percentage of patrolling and classical monocyte/macrophages in peripheral blood and spleen from indicated mouse strains (n=5). **G**, The amount of stored Guo and dGuo in patrolling and classical monocyte/macrophages in spleen from *Slc29a3*^−/−^ mice or in CD11b^+^ Ly6G^−^ NK1.1^−^ IA/IE^low^ monocyte/macrophages in wild-type spleen. \**p*<0.1, \*\**p*<0.01, \*\*\**p*<0.001.

Although accumulation of Guo and dGuo in increased patrolling monocyte/macrophages in spleen suggests that TLR7 drives histiocytosis in a cell-autonomous manner (Fig. 1G), it was possible that TLR7-dependent cytokine production promotes histiocytosis. So, we determined cytokine levels in sera of *Slc29a3^−/−^* mice. Consistent with a previous report (*16*), proinflammatory cytokines were not induced in *Slc29a3*^−/−^ mice. Serum levels of proinflammatory cytokines such as IFN-α, IFN-β, IFN-γ, IL-1β, IL-6, IL-17A, IL-23, and TNF-α were not detectable as judged by antibody arrays and/or ELISA (Fig. S5A-C). TLR7-dependent increases in serum IL-12 p40 and CXCL13 in *Slc29a3*^−/−^ mice were confirmed in ELISA (Fig. S5D, E). Despite the elevated level of IL-12 p40, its active form, IL-12 p70, were not altered (Fig. S5F). Although TLR7-mediated activation can result in inflammatory cytokine production, this was not the case with TLR7 activation by nucleosides in *Slc29a3^−/−^* mice.

Patrolling monocytes are known to clear damaged cells (*17*). So, we labelled and injected dying thymocytes to wild-type and *Slc29a3*^−/−^ mice, and found that nucleoside-laden patrolling monocytes/macrophages in the spleen of *Slc29a3*^−/−^ mice had phagocytic activity (Fig. 2A). Furthermore, professional phagocytes such as thioglycollate-elicited peritoneal exudate cells (PECs) and bone marrow-derived macrophages (BM-MCs) stored nucleosides (Fig. S6A, B), probably because they engulfed cell corpses during induction with thioglycollate *in vivo* or M-CSF *in vitro*. In contrast, less phagocytic cells such as B cells and bone marrow-derived pDCs had much less nucleoside storage (Fig. S7). Of note, dG storage was hard to detect in these poorly phagocytic cells. These results suggest that phagocytosis promotes nucleoside storage in patrolling monocytes/macrophages.

**Fig. 2.**
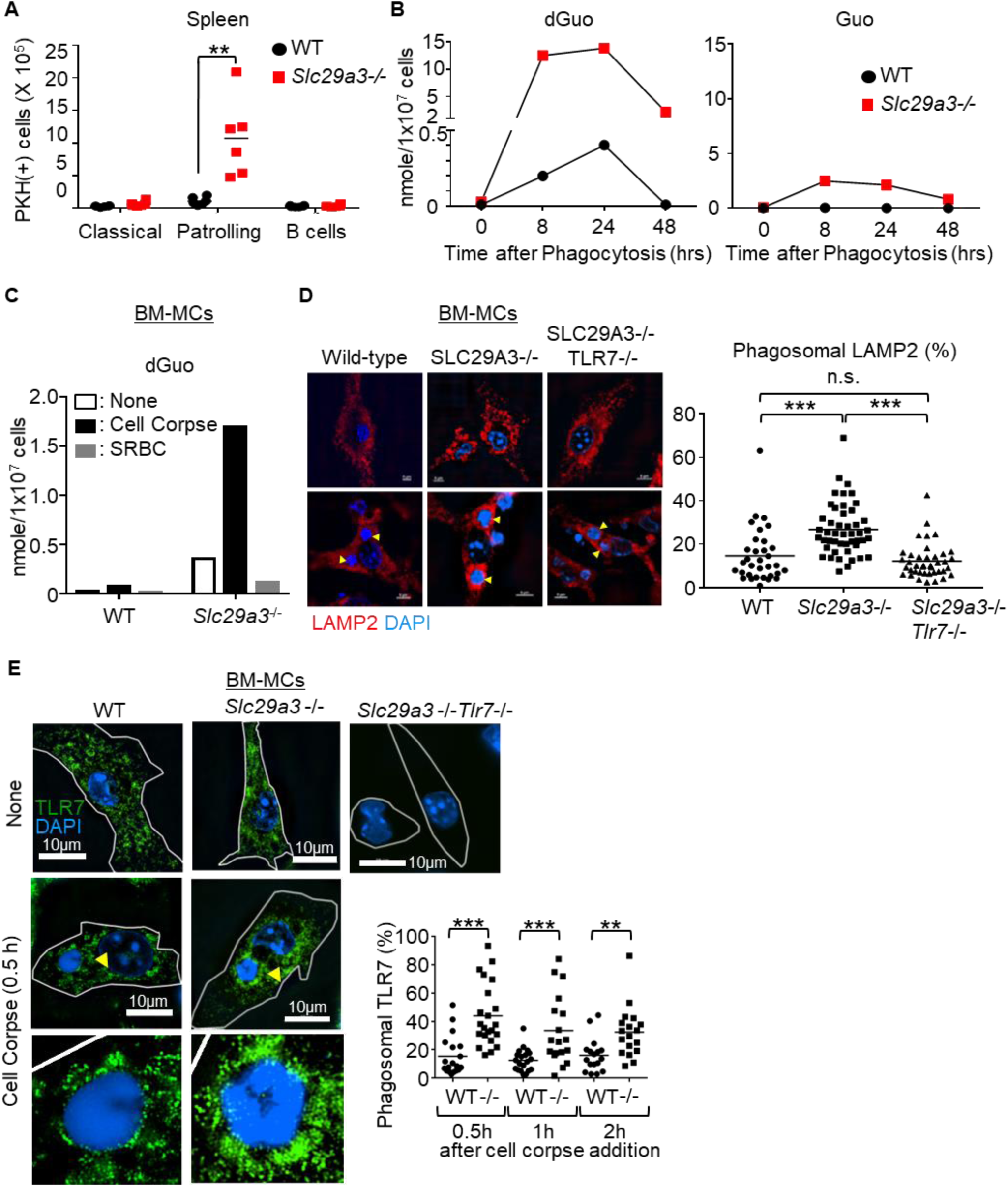
Phagocytosis-dependent nucleoside storage and TLR7 activation in *Slc29a3*^−/−^ monocyte/macrophages. **A**, The numbers of PKH26-positive immune cells in spleens at 2 hours after i.v. administration of 5 x 10^7^ PKH26-labeled dying thymocytes. **B**, **C**, The accumulation of dGuo in 10^7^ WT and *Slc29a3*^−/−^ BM-derived macrophages at 8, 24, 48 h (**B**) or 48 h (**C**) after treatment with 10^8^ dying thymocytes or 10^9^ SRBC. **D**, **E**, WT, *Slc29a3*^−/−^ and *Slc29a3*^−/−^*Tlr7*^−/−^ BM-MCs were allowed to engulf cell corpses (dying thymocytes) for 0.5 hour. Lysosomes, TLR7, and engulfed cell corpses were stained by anti-LAMP2 mAb, anti-TLR7 mAb and DAPI, respectively. Yellow arrow heads indicate phagosomes containing cell corpses. High power fields of the phagosomes indicated by yellow arrow heads are also shown (**E**). Dot plots show the percentages of LAMP2 (**D**) or TLR7 (**E**) in phagosomes. scale bar, 10μm, \*\**p*<0.01, \*\*\**p*<0.001.

To further study phagocytosis-mediated nucleoside storage, we exposed BM-MCs to dying thymocytes (Fig. 2B, Fig. S8). Accumulation of nucleosides was detected as early as at 8 h and peaked at 24 h after treatment with cell corpses. At 48 h, the levels of Guo and dGuo were decreased to steady state in wild-type BM-MCs, but remained high in *Slc29a3*^−/−^ BM-MCs. Very interestingly, dG storage was detected even in wild-type BM-MCs, suggesting that an excessive number of cell corpses can induce transient dG accumulation in lysosomes, despite SLC29A3-mediated clearance. TLR7 activation by nucleosides might occur in massive tissue damages even in wild-type monocytes/macrophages. In contrast to cell corpse phagocytosis, SRBC engulfment did not increase nucleosides (Fig. 2C, Fig. S9). Given that SRBCs do not have nuclei, nuclear DNA from cell corpses might be a major source of accumulated nucleosides. Consistent with this, DNA-derived nucleosides such as dGuo, deoxyadenosine (dAdo), deoxycytidine (dCyt), and thymidine (Thy) were accumulated only after cell corpse phagocytosis and only in *Slc29a3*^−/−^ BM-MCs (Fig. 2C, Fig. S9). Guo was not apparently increased after cell corpse phagocytosis. Stored Guo might be of endogenous origin.

As cell corpse engulfment induced nucleoside storage, we next focused on phagosomes in *Slc29a3*^−/−^ macrophages. Phagosomes containing cell corpses seemed to be similarly formed in *Slc29a3*^−/−^ BM-MCs (Fig. S10, arrowhead). Recruitment of LAMP-2^+^ lysosomes to cell corpses was found to be increased in *Slc29a3*^−/−^ macrophages (Fig. 2D), suggesting that phagolysosomal fusion was enhanced in a TLR7-dependent manner in *Slc29a3*^−/−^ mice. Because TLR7 in steady state BM-MCs was localized in LAMP2^+^ lysosomes (Fig. S11), TLR7 movement to dying cell-containing phagosomes was also increased in *Slc29a3*^−/−^ BM-MCs (Fig. 2E). Given that TLR activation increases phagolysosomal fusion in phagosome-autonomous manner (*18*), our results suggest that TLR7 in the phagosomes of *Slc29a3*^−/−^ BM-MCs was aberrantly activated, leading to additional phagolysosomal fusion. As TLR7 recruitment was lower in wild-type monocyte/macrophages, aberrant TLR7 activation by cell corpses in phagosomes would not have occurred, suggesting that SLC29A3 is localized in TLR7-containing phagosomes.

We next focused on molecular mechanisms by which TLR7 drives histiocytosis. We studied *in vitro* survival of splenic monocyte/macrophages. CD11b^+^ spleen cells from wild-type, *Slc29a3*^−/−^, and *Slc29a3*^−/−^ *Tlr7*^−/−^ mice failed to survive *in vitro* culture (Fig. 3A). In the presence of M-CSF at 1-5 ng/ml, concentrations comparable to serum M-CSF (Fig. 3B), *Slc29a3*^−/−^ CD11b^+^ cells survived *in vitro* culture, but not wild-type or *Slc29a3*^−/−^ *Tlr7*^−/−^ CD11b^+^ cells (Fig. 3A). These results suggest that TLR7-dependent hyper-responsiveness to M-CSF drives histiocytosis in *Slc29a3^−/−^* mice. Serum levels of M-CSF and CD115 expression have been reported to be increased in *Slc29a3^−/−^* mice (*16*). These increases in our *Slc29a3^−/−^* mice were dependent on TLR7 (Fig. 3B, 3C).

**Fig. 3.**
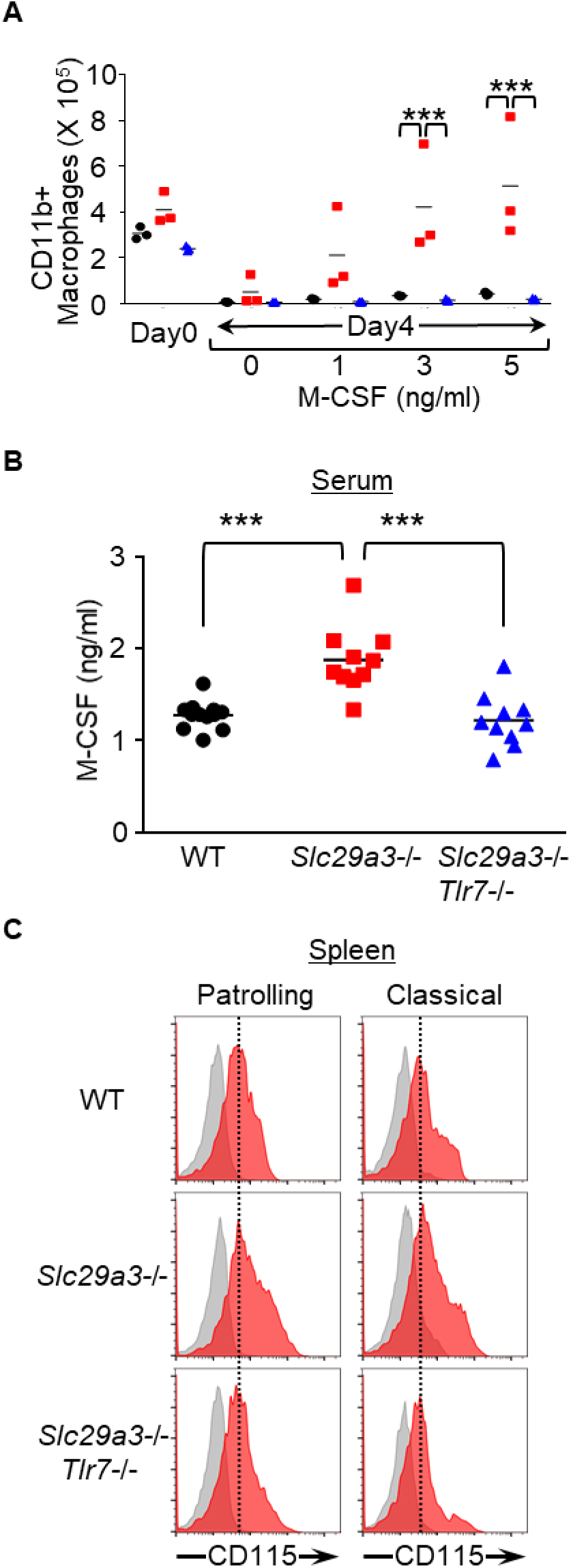
TLR7 enhances M-CSF-dependent cell survival/proliferation. **A**, The numbers of CD11b^+^ spleen cells having survived *in vitro* culture for 4 days with or without indicated concentrations of M-CSF. n=3. **B,** Serum levels of M-CSF determined by ELISA. All data were from 4∼6 months old mice (n=10). \*\*\**p*<0.001. **C**, Red histograms show expression of CD115 on splenic macrophage subsets from WT, *Slc29a3*^−/−^, and *Slc29a3*^−/−^*Tlr7*^−/−^ mice. Gray histograms represent unstained cells.

We here show that SLC29A3 negatively regulates TLR7 activation by transporting nucleosides from lysosomes to the cytoplasm (Fig. S12). When the transporting activity of SLC29A3 is impaired, stored nucleosides drive histiocytosis by promoting M-CSF-dependent survival and proliferation of monocyte/macrophages in a TLR7-dependent manner. Taken together with previous reports showing that TLR7 and TLR8 bind and respond to nucleosides (*9, 11*), SLC29A3 disorders in mice are caused by TLR7 responses to nucleosides.

TLR7 was localized in LAMP-2^+^ compartment and recruited to phagosomes containing cell corpses due to constitutive lysosomal fusion. Very importantly, lysosomal fusion was increased in *Slc29a3*^−/−^ macrophages in a TLR7-dependent manner. Given that phagolysosomal fusion is activated by phagosomal TLRs in phagosome-autonomous manner (*19*), phagosomal TLR7 is likely to be activated to promote further lysosomal fusion in *Slc29a3*^−/−^ macrophages. In addition to phagosomal maturation, TLR7 activation in phagosomes is likely to drive histiocytosis. Most importantly, TLR7 responses to nucleosides in *Slc29a3*^−/−^ mice drove cell proliferation/survival without any apparent inflammation. TLR7 might activate two distinct types of responses, proinflammatory and pro-histiocytosis responses. Monocytosis with thrombocytopenia is reported in mice where TLR7 is constitutively activated by its over-expression or altered trafficking (*5, 6*). These phenotypes are understood as macrophage activation syndrome (MAS), a complication of autoimmune diseases and infections (*8*). TLR7-dependent pro-histiocytosis responses might contribute to MAS.

In conclusion, we here show that TLR7 drives histiocytosis in SLC29A3 disorders. Further study needs to focus on the role of TLR7 in histiocytosis caused by viral infection or autoimmune diseases. TLR7 might be a target for therapeutic intervention in histiocytosis.

## Acknowledgments

We thank: Prof. P. W. Kincade for critically reviewing the manuscript; Noriko Tokai for helping us analyze the imaging results; the IMSUT FACS Core laboratory for assistance with the flow cytometric analysis and the cell sorting. This work was supported in part by: Grant-in-Aid for Scientific Research (S) to K. M. (16H06388), (A) to T. Shimizu. (16H02494), (B) to S.-I.S. (26293083), (B)(19H03164) to U.O, and (C) (16K08827) /(B) (19H03451) to T.S.; Grant-in-Aid for Scientific Research on Innovative Areas (18H04669,19H04950) to U.O., (18H04856) to S.-I.S., and (18H04666) to K.M. ; Grant-in-Aid for Young Scientists (19K16685) to R.S. ; CREST to T.S.,T.I. and T. Shimizu.; AMED (JP18ek0109385) to T.S.; and Joint Research Project of the Institute of Medical Science at the University of Tokyo. T.S. and K.M. conceived and designed the experiments. T.S. and N.Y. constructed knock-out mice with the help of M. K. and C. K.. T.S. performed most of the experiments except LC-MS Analysis, High-Resolution Microscopy and Electron Microscopy. M.T., Y.Y. and T.I. performed LC-MS analysis. High-Resolution Microscopy and Electron Microscopy were carried out by S.-I.S. and H.S., respectively. T.S. and K.M. wrote the paper with assistance from M.T., S.-I.S., H.S., R.F., R.S., K.F., Y.M., U.O., and T. Shimizu. K.M. and T.S acquired funding for conducting this research project. All authors reviewed and edited the results and comments on the manuscript.

## Methods

### Generation of *Slc29a3*^−/−^ and *Tlr7*^−/−^ Mice

SLC29A3 Knock-out (*Slc29a3*^−/−^) and TLR7 Knock-out (*Tlr7^−/−^*) mice were generated using a CRISPR/CAS system. gRNA target sites of *Slc29a3* and *Tlr7* genes were determined by CRISPRdirect software (http://crispr.dbcls.jp/). 5’-agcttcttgatggttactcg-3’ and 5’-gaacagttggccaatctctc-3’ sequences were used for construction of *Slc29a3*^−/−^ and *Tlr7^−/−^* mice, respectively. These gRNA target sequences were cloned into the BbsI site of pKLV-U6 gRNA (BbsI)-PGKpuro2ABFP Vector (Addgene plasmid 50946), and gRNAs were synthesized from the constructed vector *in vitro* using MEGAshortscript T7 Transcription Kit (ThermoFisher Scientific, MA, USA). In addition, hCAS9 mRNA was synthesized from the hCAS9 sequence in pX458 (Addgene plasmid 48138) *in vitro* using mMESSAGE mMACHIN T7 ULTRA Transcription Kit (ThermoFisher Scientific). To construct *Slc29a3*^−/−^*Tlr7^−/−^* double knock-out mice, gRNAs (50ng/μl) to target *Slc29a3* and *Tlr7^−^* genes, hCAS9 mRNA(100ng/μl), and the single-stranded DNA (100ng/μl) to introduce a stop codon into the *Slc29a3* locus (5’-tgcctctgaggacaatgtataccacagctccaatgctgtctacagagcccTGATAGCGTAAAGCACTGAGGAAGcgagtaaccatcaagaagctgaccaggaagccctgctggggaaactacta-3’ ; Lower-case letters and capital letters represent Homology Arm and knock-in sequences, respectively) were injected into zygotes from C57BL/6 mice at the pronuclei stage. The injected zygotes were transferred to the oviducts of pseudopregnant female C57BL/6 mice. Candidates of *Slc29a3*^−/−^ *Tlr7^−/−^* double knock-out were typed by PCR using primer pairs for the SLC29A3 gene (Fw: 5’-CCAGCATGGACGAGAGATGTCTTC-3 ’, Rv: 5’-GCACCATTGAAGCGATCCTCTGG-3’) and the TLR7 gene (Fw: 5’-GAGGGTATGCCGCCAAATCTAAAGAATC-3 ’, Rv: 5’-CTGATGTCTAGATAGCGCAATTGC-3’). The PCR product was analyzed by a MCE-202 MultiNA Microchip Electrophoresis System for DNA/RNA Analysis (SHIMAZU, Tokyo, Japan), and one candidate male mouse having both knock-in alleles in SLC29A3 gene and a frame shift mutation in TLR7 gene was chosen for mating with wild-type B6 female mice. Obtained (wild-type B6/dKO) F1 mice were used for the construction of both *Slc29a3*^−/−^ and *Slc29a3*^−/−^ *Tlr7^−/−^* mice. The sequences of mutation allele in *Slc29a3*^−/−^ and *Tlr7^−/−^* mice was confirmed by the direct sequencing performed by FASMAC (Kanagawa, Japan).

### Mice

Wild-type C57BL/6 mice were purchased from Japan SLC, Inc. (Shizuoka, Japan). All animals were housed in specific pathogen-free facilities at the Institute of Medical Science, University or Tokyo (IMSUT). All animal experiments were approved by the Institutional Animal Care and Use Committee in the IMSUT.

### Reagents

Guanosine (Guo) and Uridine (Uri) were purchased from Wako Pure Chemical Industries (Osaka, Japan). Adenosine (Ado), Cytidine (Cyt) and Thymidine (Thy) were purchased from MP Biomedicals (Santa Ana, CA, USA). 2’-Deoxyadenosine (dAdo) and 2’-Deoxycytidine (dCyt) were purchased from Sigma-Aldrich Japan (Tokyo, Japan). 2’-Deoxyuridine (dU) and 2’-Deoxyguanosine monohydrate (dG) were purchased from Jena Bioscience (Jena, Germany).

Stable isotope labeled nucleosides, G (13C10, 98%;15N5,96-98%), U(13C9, 98%;15N2,96-98%), A(13C10, 98%;15N5,96-98%), C(13C9, 98%;15N3,96-98%) and dG(13C10, 98%;15N5,96-98%) were purchased from Cambridge Isotope Laboratories (Massachusetts, USA)

### Proteome Profiler

Relative expression levels of 111 plasma cytokines were quantified by the Proteome Profiler Mouse XL Cytokine Array (R&D Systems). The antibody array was performed according to the manufacturer’s instruction. To achieve maximum sensitivity, 200μl plasma was used as sample and antibody array membrane was incubated with plasma samples overnight at 4°C. After overnight incubation, membranes were sequentially reacted with Detection Antibody Cocktail and Streptavidin-HRP. Finally, membranes were treated with ECL Select Western blotting detection Reagent (GE Healthcare). The chemiluminescent signals on membranes were quantified by ImageQuant LAS 500 imager system (GE Healthcare), and the intensity of each spot was quantified using ImageJ software (National Institutes of Health). In addition, serum concentrations of several cytokines (IL-1β, TNF-α, IL-6, IL-12p40, CXCL13 and M-CSF) were further determined by ELISA to confirm the results from Proteome profiler cytokine array.

### Flow Cytometry and Cell Sorting

Peripheral blood mononuclear cells (PBMCs), bone marrow cells, and spleen cells were stained with fluorescent dye-conjugated monoclonal antibodies specific for the following markers: CD11b (clone M1/70), FcγRIV (clone 9E9), CD3ε (clone 145-2c11), CD19 (clone 6D5), CD11c (clone N418), CD71 (clone R17217), Ter119/Erythroid Cells (clone Ter119), NK1.1 (clone PK136), Ly6C (clone HK1.4) and Ly6G (clone 1A8). These antibodies were purchased from eBioscience (San Diego, CA, USA), Biolegend (San Diego, CA, USA), BD Biosciences (San Jose, CA, USA) or TONBO biosciences (San Diego, CA, USA). Biotinylated anti-mouse TLR7 (A94B10) mAb was previously established in our laboratory^20^.

For the preparation of single cell suspension, spleens were minced by slide glasses and bone marrow cells were pipetted several times to disperse the cells in RPMI1640 medium. Suspended samples were teased through nylon mesh to remove tissue debris. All samples were treated with BD Pharm Lyse lysing buffer (BD Biosciences, Tokyo, Japan) to remove red blood cells before subjecting to cell staining. Cell staining for flow cytometry analysis was performed in staining buffer (1×PBS with 2.5% FBS and 0.1% NaN3). Single cell suspension from mice was incubated for 10 minutes with anti-CD16/32 (clone 95) blocking antibody diluted in staining buffer. Then cells were stained by fluorescein-conjugated mAbs for 15 minutes on ice. To detect endolysosomal TLR7, cells after cell surface staining were fixed and permeabilized by Fixation/Permeabilization Solution Kit (BD Biosciences), and stained again by biotinylated anti-mouse TLR7 mAb and PE streptavidin (Biolegend, USA). Stained cells were analyzed by the BD LSR Fortessa cell analyzer (BD Biosciences).

Bone marrow derived pDCs for sorting were stained in sorting buffer (1×PBS with 10% FBS, 10mM HEPES and 1mM Sodium Pyruvate). Mouse splenic immune cells for sorting were stained in staining buffer. All cells were sorted by the FACSAria flow cytometer (BD Biosciences).

### Platelet and Leukocyte Count

Platelet, leukocyte number in PBMC was analyzed by automatic hematology analyzer, Celltac α (Nihon Kohden, Tokyo). The numbers of spleen and bone marrow cells were determined by a Countess II FL Automated Cell Counter (Thermo Fisher Scientific, Tokyo, JAPAN).

### Preparation of Mouse Bone Marrow (BM) Derived Macrophages, cDCs and pDCs

BM cells were collected from tibiae, femora and pelves of wild-type, *Slc29a3*^−/−^, and *Slc29a3*^−/−^*Tlr7*^−/−^ mice, and red blood cells were removed by BD Pharm Lyse lysing buffer treatment. For preparation of BM-macrophages (BM-MCs), BM cells were plated at a density of 7 x10^6^ cells per one non-tissue culture polystyrene 94mm petri dish (Greiner Bio-One, Frickenhausen, Germany) and cultured in 10ml RPMI medium (Gibco, Paisley, UK) supplemented with 10% FBS, Penicillin-Streptomycin-Glutamine (Gibco, Paisley, UK), 50μM 2-ME and 100ng/ml recombinant murine macrophage colony stimulating factor (M-CSF, PeproTech Inc., Rocky Hill, NJ, USA) for 6 days. For BM-cDCs, BM cells were plated at a density of 1×10^6^ cells per well in 24-well plates and cultured in 1ml RPMI 1640 medium (Gibco, Paisley, UK) supplemented with 10% FBS, Penicillin-Streptomycin-Glutamine, 50μM 2-ME and 10 ng/ml of recombinant murine granulocyte– macrophage colonystimulating factor (GM-CSF, PeproTech Inc., Rocky Hill, NJ, USA) for 7 days. Half volume of medium was changed every other day. For BM-pDCs, BM cells were plated at a density of 2.5×10^7^ cells per one 10cm cell culture dishes (Greiner Bio-One, Frickenhausen, Germany) and cultured in 10ml RPMI 1640 medium (Gibco, Paisley, UK) supplemented with 10% FBS, Penicillin-Streptomycin-Glutamine, 50μM 2-ME and 100 ng/ml of recombinant murine fms-like tyrosine kinase-3 ligand (Flt3L, PeproTech Inc., Rocky Hill, NJ, USA) for 7days. Flt3L induced pDCs were then stained by anti-CD11c/B220 mAbs, and CD11c^+^B220^+^ cells were sorted by FACSAria flow cytometers (BD Biosciences, San Jose, CA, USA) as BM-pDCs and subjected to experiments.

### Cytokines Measurement by ELISA

Serum cytokine mice was measured by ELISA. Blood samples were allowed to clot for 1hours at room temperature before centrifugation for 10 minutes at 2000g. Collected serum was subjected to ELISA to determine the concentration of Mouse IL-12p40, TNF-α, IL-6 and IL-1β by Ready-Set-Go! ELISA kits (eBioscience, CA, USA), Mouse M-CSF and CXCL13 by Quantikine ELISA Kit (R&D Systems, MN, USA), and mouse IFN-α and IFN-β by VeriKine ELISA Kit (PBL Assay Science, NJ, USA).

### Cell Death Induction and Phagocytosis Assay

Cell death was induced by treatment of thymocytes or splenocytes at 47 °C for 20 minutes, and then incubated at 37 °C for 3 hours before subjecting to phagocytosis assay. For phagocytosis assay in vivo, dead cells were stained by PKH26 Red Fluorescent Cell Linker Kit for General Cell Membrane Labeling (Sigma-Aldrich, MO, USA) according to manufacturer’s instructions. 5 × 10^7^ PKH26 stained dead cells were then intravenously administered to mice and spleens were collected at 0.5 or 2 hours after dead cells administration. Immune cells engulfing PKH26 positive dead cells were analyzed by flow cytometry.

### Lentiviral Transduction

Lentiviral transduction was used to overexpress gRNA and mRNA in macrophage cell line J774. Constructs were subcloned into the pKLV-U6gRNA(BbsI)-PGKpuro2ABFP vector (Addgene plasmid 50946) and ViraPowe Lentiviral expression system (Thermo Fisher Scientific, USA) was used to prepare lentivirus according to the manufacturer’s instructions. Supernatants containing lentivirus were directly added to the wells culturing J774 cells for transduction.

### Structured Illumination Microscopy

wild-type, *Slc29a3*^−/−^, and *Slc29a3*^−/−^ *Tlr7*^−/−^ BM-macrophages were allowed to adhere to collagen-coated coverslips overnight. Macrophage cell lines J774, cells were allowed to adhere to collagen-coated coverslips overnight and activated with 1μM Phorbol 12-myristate 13-acetate (PMA) for 2h. Attached cells on coverslips were treated with the heat treated dying thymocytes for 1h. After engulfment, cells were fixed with 4% paraformaldehyde for 10min, permeabilized with 0.2% Saponin in PBS for 30min, and blocked with 2.5% BSA Blocking One (Nacalai tesque, Japan) for 30min. The cells were then incubated with anti-TLR7 antibody and anti-HA antibody (Roche, USA) at 37°C for 90min, and washed three times, then incubated for 90 min at 37°C with goat anti-mouse and Rat antibodies conjugated to AlexaFluor-488 or 568 and DAPI (Invitrogen). All microscopy was performed using Nikon Structured illumination microscopy (N-SIM) at excitation wavelengths of 405nm, 488nm, 541nm with100xH NA1.49 CFI Apochromat TIRF (N-SIM, Nikon). Data Acquisition was performed in 3D SIM mode before image reconstruction in NIS-Element software. The TLR7 fluorescence intensity in phagosomal compartment was then calculated in NIS-Element. Phagosomal TLR7 (%) was calculated by quantitating the fluorescence intensity with phagosomal TLR7 and total cellular TLR7. Each image is representative of at least three independent experiments. Statistical significance was determined using two-sided t tests.

### Electron Microscopy

5 × 10^7^ Heat killed thymocytes were added to 5 × 10^6^ mouse BM-macrophages (BM-MCs) in 10cm dishes and cultured for 1hour. Then BM-MCs on plates were fixed in the solution containing 2.5% glutaraldehyde, 2% formaldehyde in 0.1M sodium phosphate buffer (pH 7.4) for 2hrs at room temperature. After fixation, tissues were rinsed and post-fixed in the 2% osmium tetroxide in the same buffer solution on ice. They were then washed, dehydrated in the graded series of ethanol and embedded in Epon 812 resin mixture (TAAB, Berks, UK). Semi-thin sections of about 0.7 micro-meter thickness were cut on a Reichert Ultracut N ultramicrotome, stained with 0.2% toluidine blue and examined under a Zeiss Axioskop microscope. Ultra-thin sections were cut, stained with uranyl acetate and lead citrate and examined with a HITACHI H-7500 electron microscope.

### LC-MS Analysis (Cell Preparation)

The quantitative nucleoside analysis was performed with a LC-MS system equipped with a reversed-phase column (2.0 mm I.D. × 100 mmL) packed with Develosil C30 UG (3μm particle, Nomura Chemical) connected to a hybrid quadrupole-Orbitrap mass spectrometer (Q Exactive, Thermo Fisher Scientific) through an electrospray interface. 1 × 10^7^ Sample cells were lysed with D solution (7M guanidine hydrochloride and 0.5M Tris-HCl/10mM EDTANa2, pH 8.5) containing stable isotope labeled nucleosides (final standard nucleoside concentration; 1 nmol/400 μL of A/U/G/C/dG). 100mg of tissues were lysed with D solution containing stable isotope labeled nucleosides (final standard nucleoside concentration; 1 nmol/400 μL of C/dG, 10nmol/400μL of U/G and 100nmol/400μL of A). The extract was centrifuged at 10,000 x g for 30 min and the supernatant was diluted 40- to 200-folds with 10mM ammonium acetate buffer (pH 6.0). Samples (ca 1∼100pmole nucleosides/40 μL) were loaded to a reversed phase column and eluted with a 30 min linear gradient from 2% to 12% acetonitrile in 10mM ammonium acetate buffer (pH 6.0) at a flow rate 100 μL/min. The eluate from the first 6 min was wasted automatically by switching a 3-way electric valve to remove guanidine hydrochloride from the system and was subsequently sprayed into a mass spectrometer at 3.0 kV operating under the positive ion mode. The mass spectra were acquired at a mass resolution of 35,000 from m/z 200 to 305. Each nucleoside in the sample cells or tissues was quantitated from the peak height relative to that of the corresponding isotope labeled standard nucleoside. All LC-MS data were processed and analysed using Xcalibur (version 3.0.63, Thermo) and Excel 2013 (Microsoft).

### Statistical Analysis

Statistical analysis to compare two conditions was performed using Two-tailed unpaired t test with Holm-Sidak correction. All data are presented as mean value ± standard deviation (s.d.). P<0.05 was considered statistically significant. *P < 0.05, **P < 0.01, ***P < 0.001.

## Supplementary Materials

**Figure S1.**
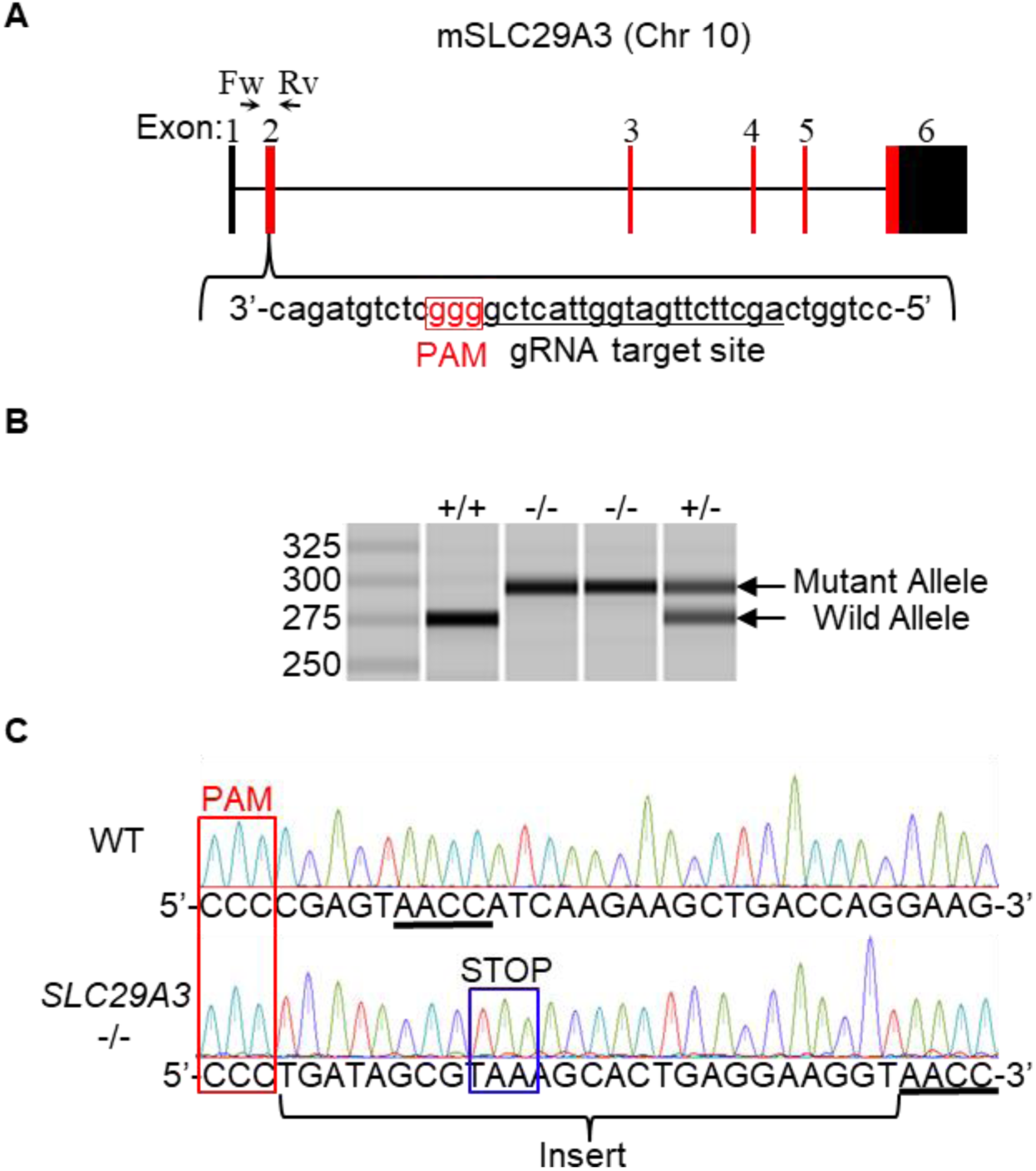
Generation of *Slc29a3*^−/−^ mice. **A**, Genomic configuration of the *SLC29A3* gene and the gRNA target site to introduce a mutation using a CRISPR/CAS system in the Exon 2 of *SLC29A3* gene. The PAM sequence is shown in the red box. **B**, Genomic PCR with the primer set (Fw and Rv) shown in (**A**) to reveal the size change of PCR products due to insertional mutation in the target site. **C**, Direct sequencing of the gRNA target site in WT and *Slc29a3*^−/−^ mice. The inserted sequence containing stop codon is indicated.

**Figure S2.**
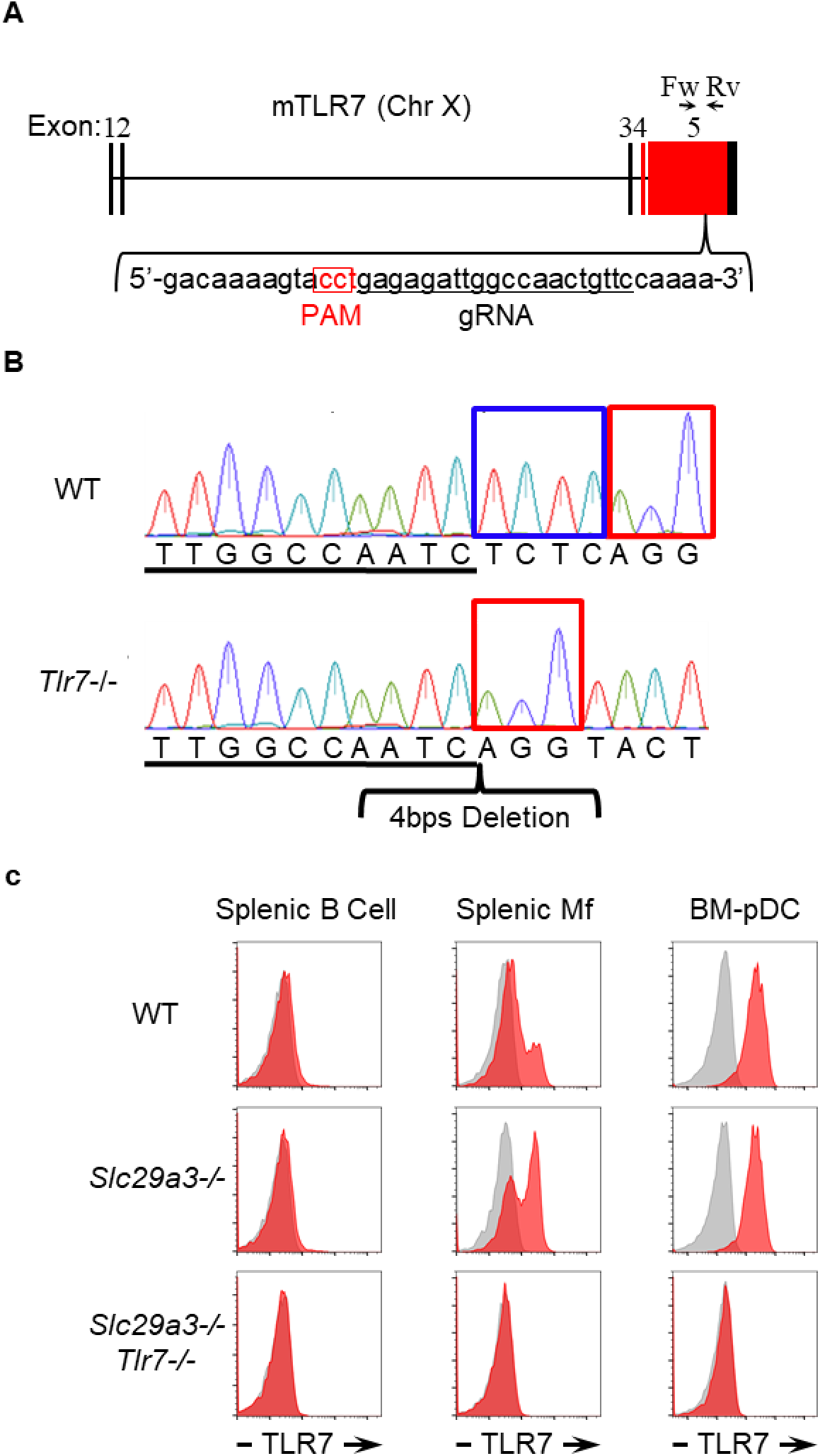
Generation of *Tlr7*^−/−^ mice. **A,** Genomic configurations of the *Tlr7* gene and the gRNA target sites to introduce a mutation using a CRISPR/CAS system in the Exon 5 of *TLR7* gene. The PAM sequence is shown in the red box. **B**, The sequence data of the gRNA target site in WT and *Tlr7*^−/−^ mice to show the 4-bp deletion mutation indicated by a blue box. The PAM sequence is shown in the red box. **C**, FACS analyses to show intracellular TLR7 expression level in splenic B cells, splenic macrophages, and BM-derived plasmacytoid DCs from WT, *Slc29a3*^−/−^, and *Slc29a3*^−/−^ *Tlr7*^−/−^ mice. Red and gray histograms show membrane-permeabilized staining with or without anti-TLR7 mAb, respectively.

**Figure S3.**
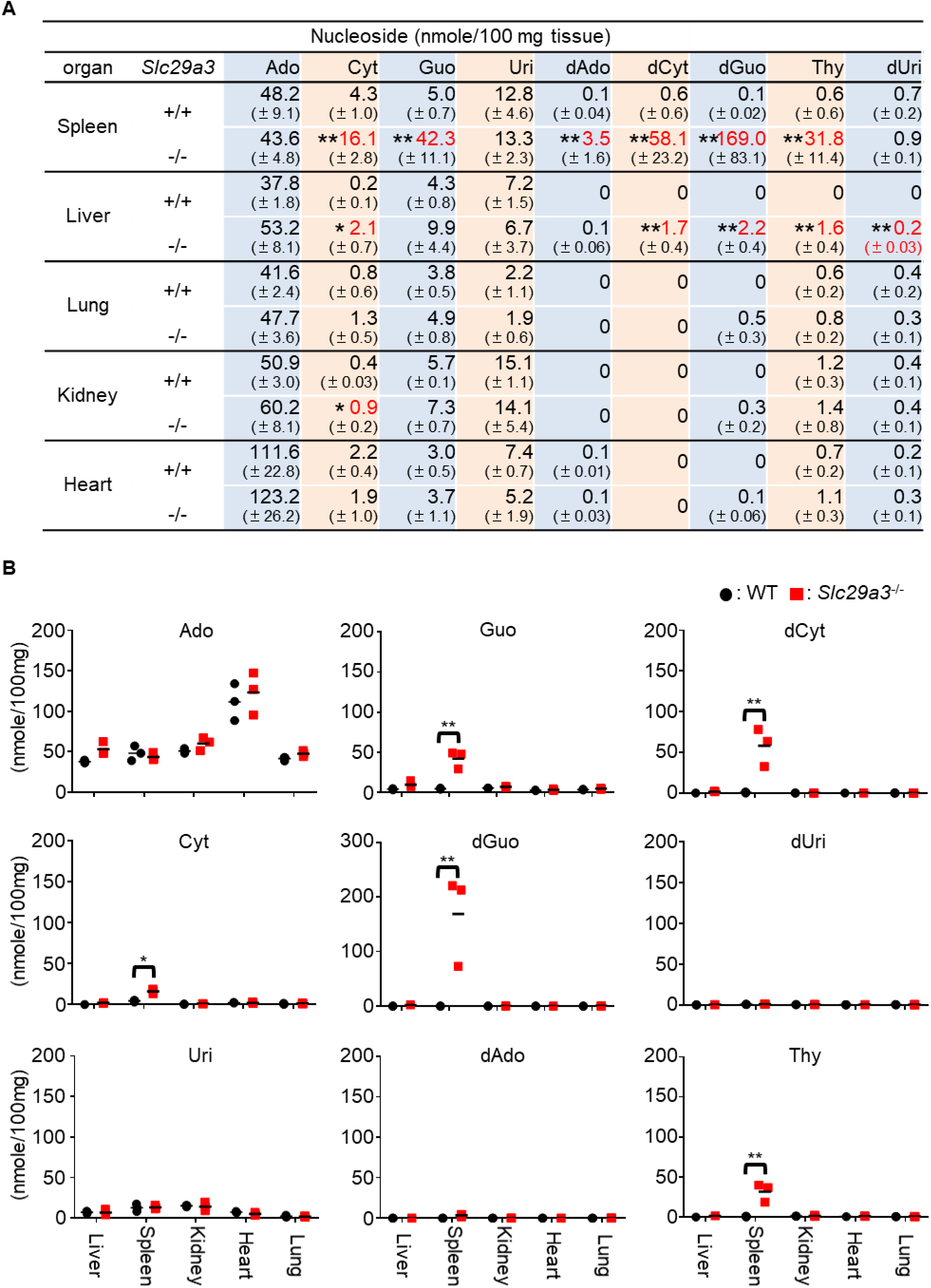
Nucleoside storage in *Slc29a3*^−/−^ mice. **A**, **B**, The amounts of nucleosides in various tissues from WT and *Slc29a3*^−/−^ mice were determined by LC-MS analyses (n=3). The results are shown as both a table (**A**) and a dot plot (**B**). The results are presented as mean values ± s.d. from three samples.

**Figure S4.**
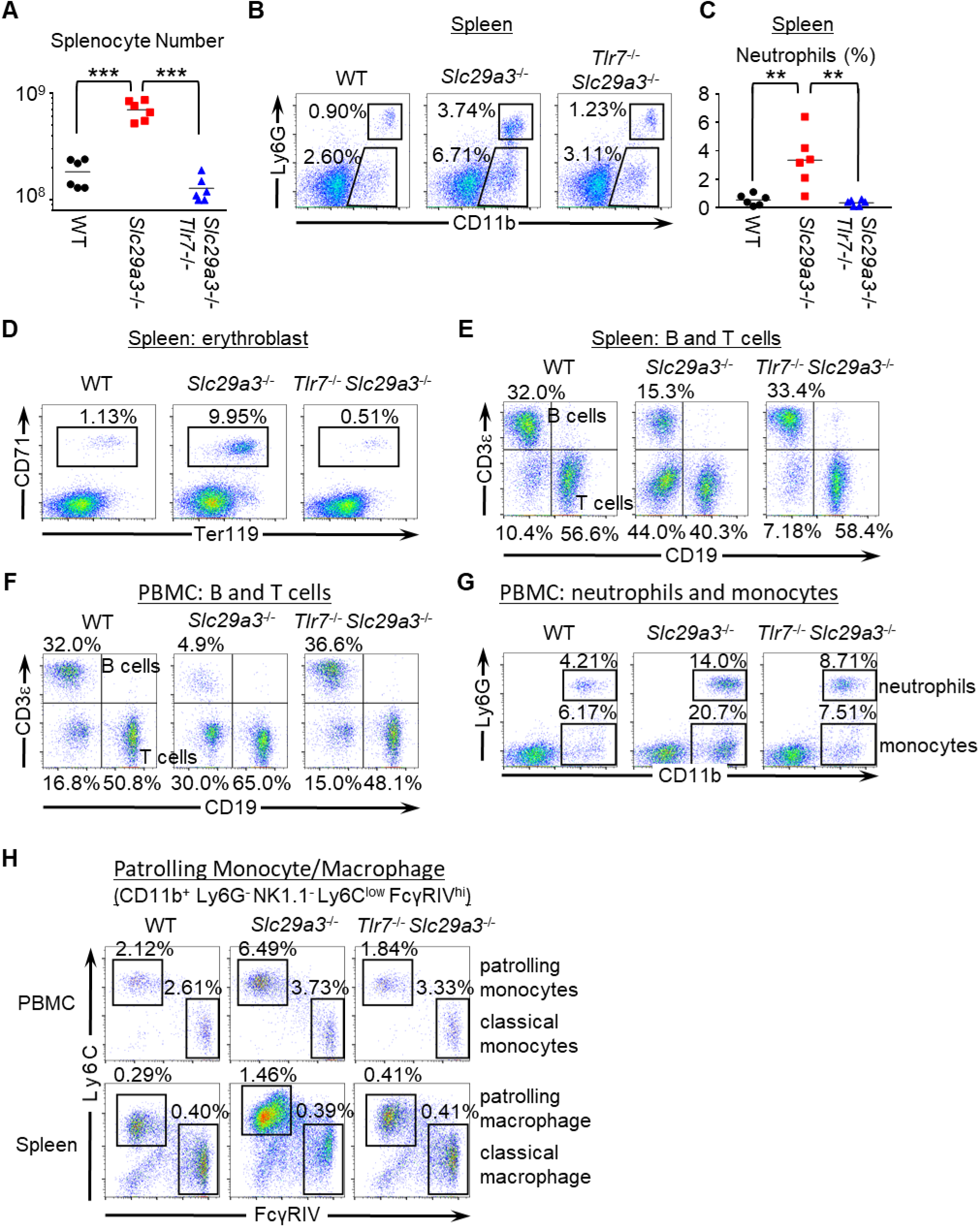
TLR7-dependent monocytosis in *Slc29a3*^−/−^ mice. Immunocytes from WT, *Slc29a3*^−/−^, and *Slc29a3*^−/−^ *Tlr7*^−/−^ mice were analyzed by flowcytometry. **A**, Dot plots show the number of spleen cells in indicated mouse strains. (n=6) **B**, FACS analyses to show expression of Ly6G and CD11b on spleen cells. Ly6G^+^ neutrophils and Ly6G^−^ monocytes/macrophages are indicated by the gates. **C**, Dot plots showing the percentages of splenic neutrophils. **D**, **E**, FACS analyses to show splenic CD71^+^ Ter119^+^ erythroblasts (**D**), CD3ε^+^ T cells (**E**), and CD19^+^ B cells (**E**). Their percentages are also shown. **F**, **G**, FACS analyses to show expression of CD3ε, CD19, Ly6G, and CD11b in peripheral blood mononuclear cells (PBMC). The percentages of CD3ε^+^ T cells (**F**), CD19^+^ B cells (**F**), Ly6G^+^ neutrophils (**G**), and Ly6G^−^ monocytes (**G**) are also shown. **H**, FACS analyses to study patrolling (Ly6C^low^ FcγRIV^hi^) and classical (Ly6C^hi^ FcγRIV^low^) monocyte/macrophages in peripheral blood and spleen from wild-type (WT), *Slc29a3*^−/−^ and *Slc29a3*^−/−^ *Tlr7*^−/−^ mice.

**Figure S5.**
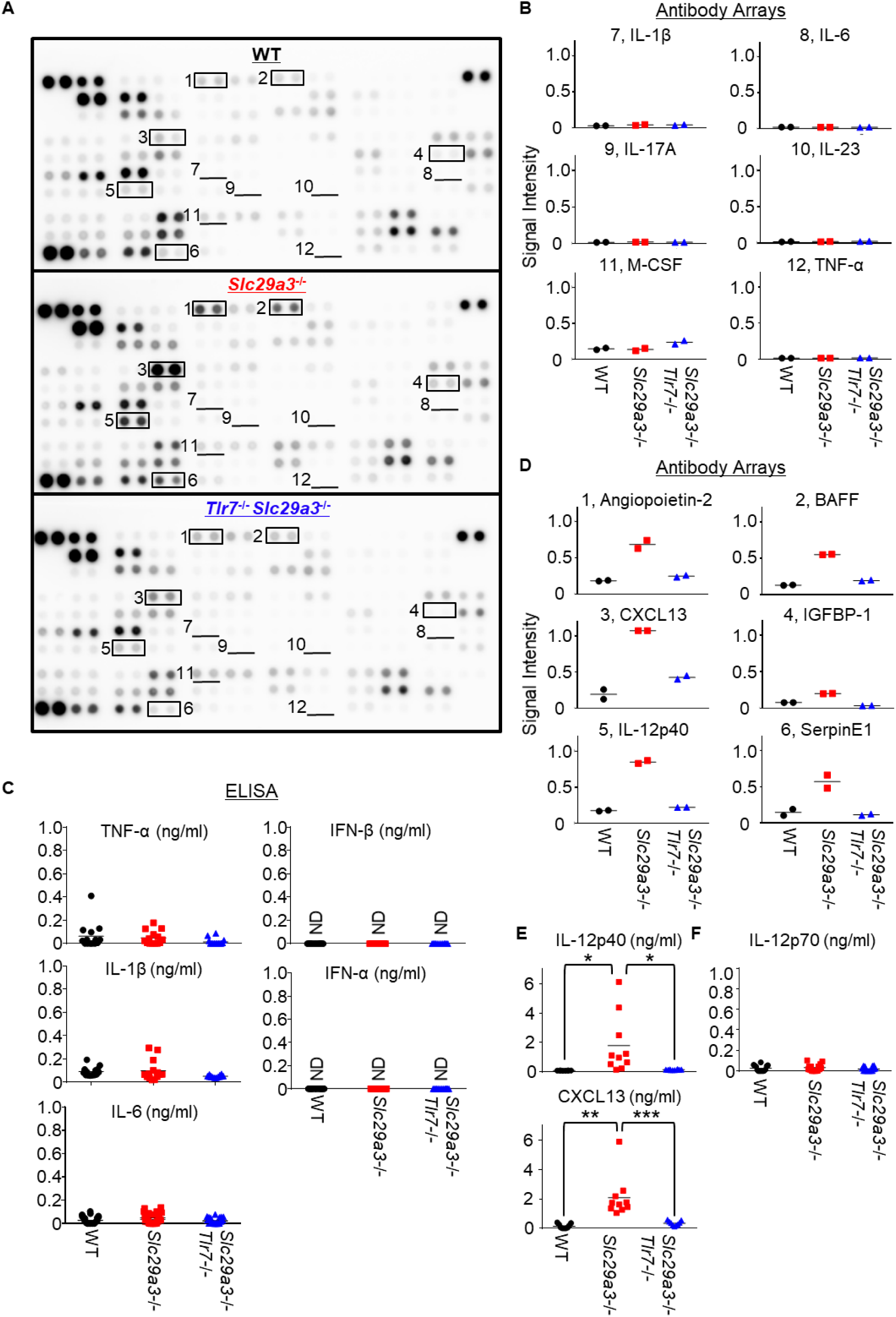
Serum cytokine levels in *Slc29a3*^−/−^ mice. **A**, The levels of 111 plasma proteins from 6 months old WT, *Slc29a3*^−/−^ and *Slc29a3*^−/−^ *TLR7*^−/−^mice, were evaluated by Proteome Profiler antibody array. **B**, Densitometry analyses of the results of the antibody array. Each dot represents the spot signal intensity of each protein divided by that of control spots from (**A**). **C**, Serum levels of TNF-α, IL-1β, IFN-β, IL-6, and IFN-α were determined by ELISA. All data was from 4∼6 months old mice (n=14). \*\*\**p*<0.001. **D**, Shown are the densitometry analyses of the plasma proteins that were increased in *Slc29a3*^−/−^ mice in a TLR7-dependent manner. Each dot represents the spot signal intensity of each protein divided by that of control spots from (**A**). **E**, **F**, Serum levels of IL-12 p40, CXCL13, and IL-12 p70 were determined by ELISA as in (**C**).

**Figure S6.**
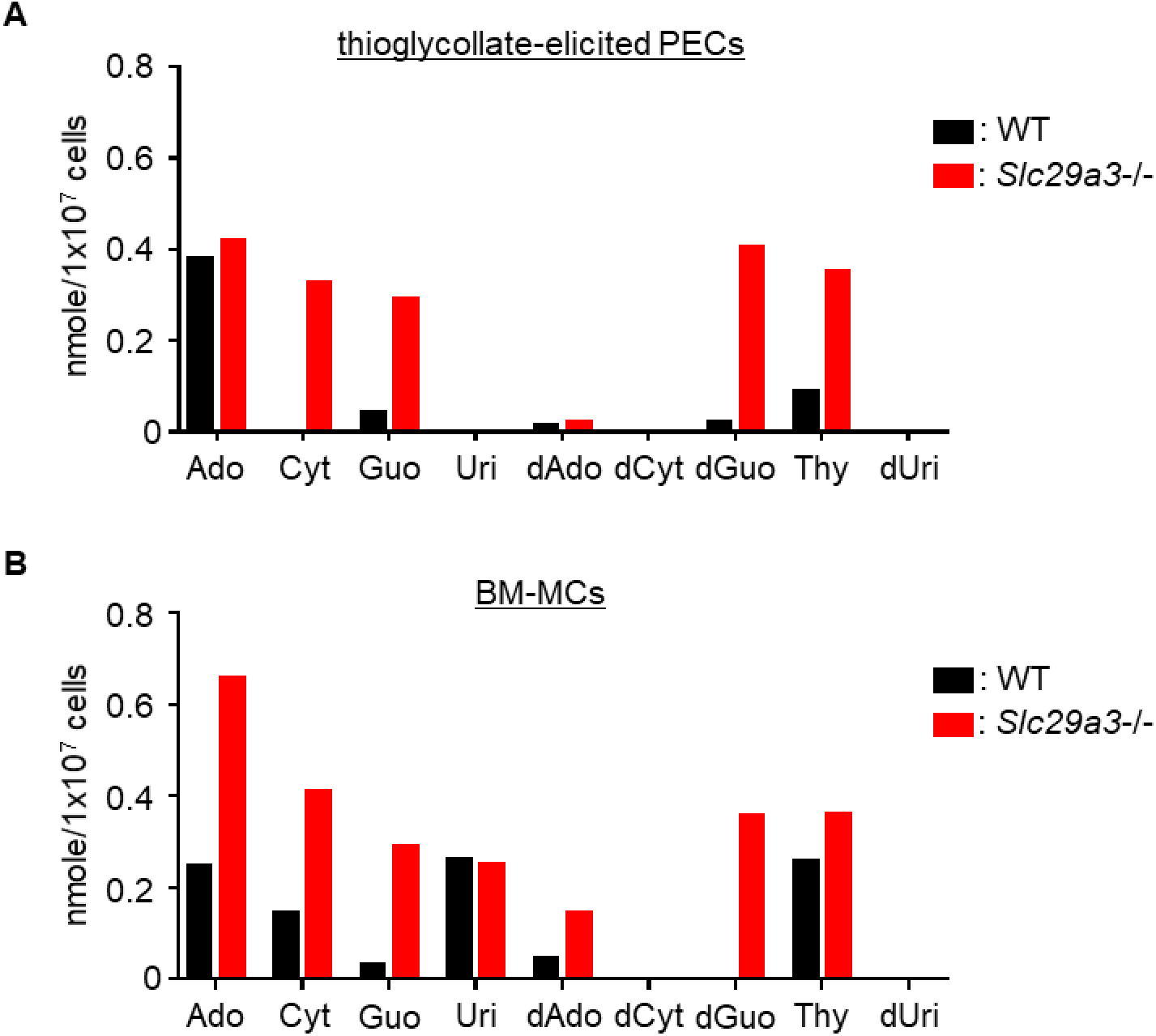
Nucleoside storage in *Slc29a3*^−/−^ macrophages. The amounts (nanomole) of accumulated nucleosides (nanomole) in 10^7^ cells of thioglycollate-elicited peritoneal exudate cells (PECs)(**A**) and bone marrow-derived macrophages (BM-MCs) (**B**) from WT (black) and *Slc29a3*^−/−^ mice (red) are analyzed by LC-MS.

**Figure S7.**
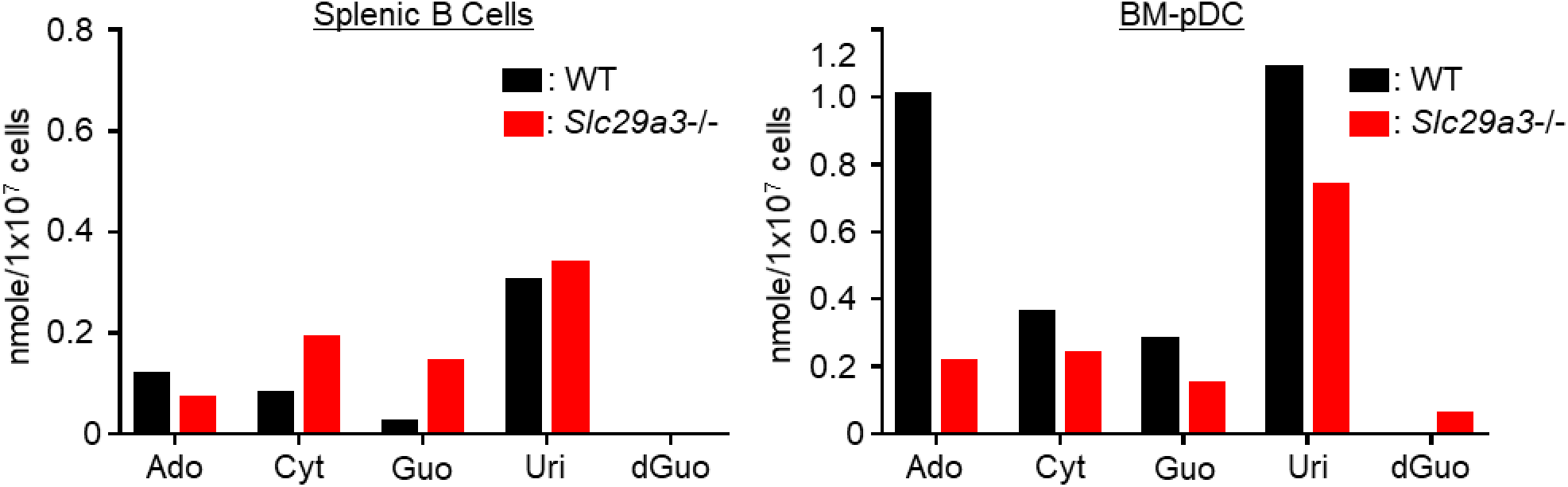
Nucleoside storage in B cells and plasmacytoid DCs. The amounts (nanomole) of accumulated nucleosides (nanomole) in 10^7^ cells of splenic B cells and bone marrow-derived plasmacytoid DCs (BM-pDCs) from WT and *Slc29a3*^−/−^ mice are analyzed by LC-MS.

**Figure S8.**
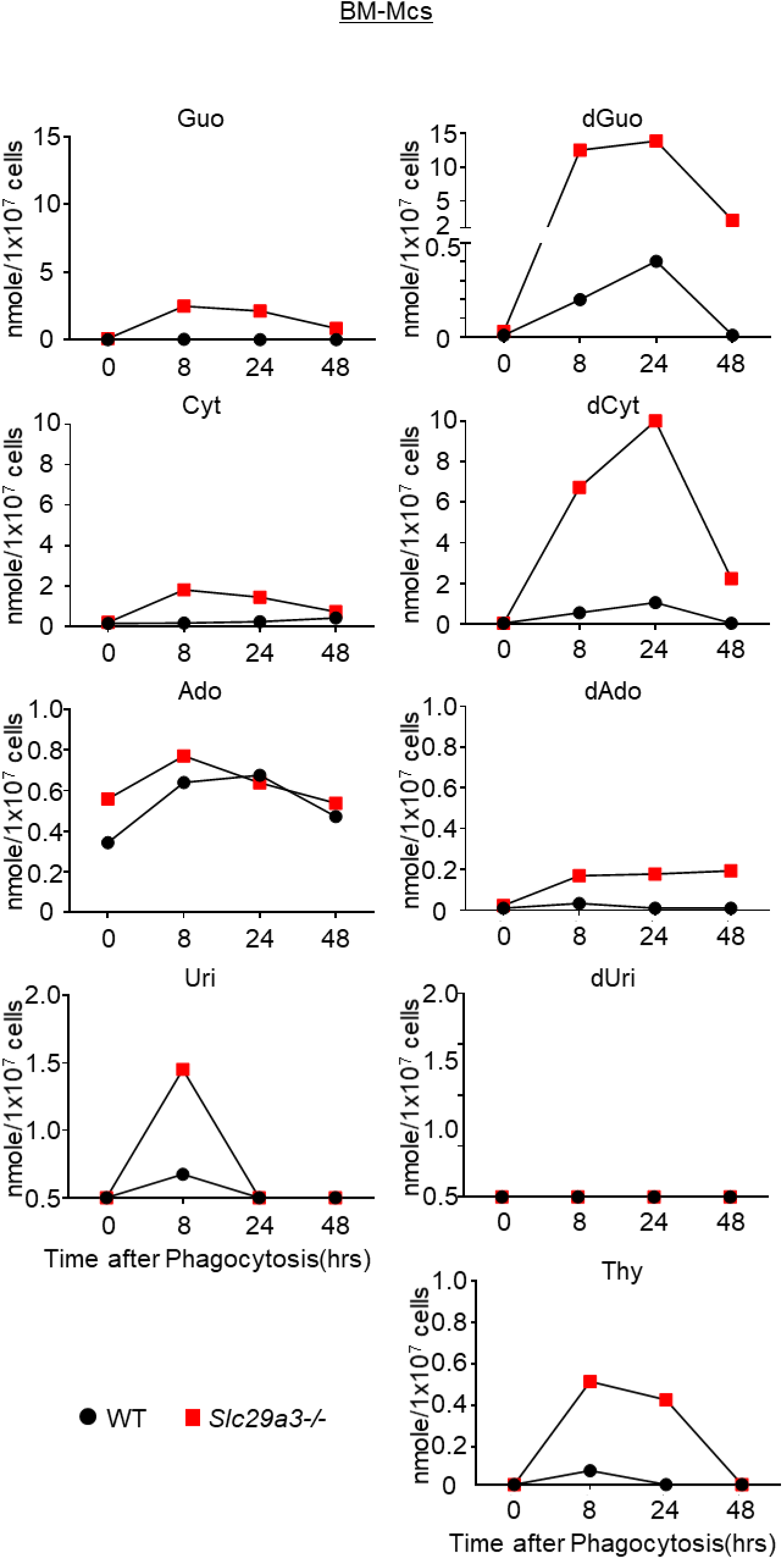
Time course of nucleoside storage in macrophages after cell corpse engulfment. Accumulation of nucleosides in 10^7^ WT (black circle) and *Slc29a3*^−/−^ (red square) BM-derived macrophages at 8, 24, 48 h after treatment with 10^8^ dying thymocytes was determined by LC-MS analyses. The data for Guo and dGuo are shown in Fig. 2B. Similar results were obtained from three independent experiments.

**Figure S9.**
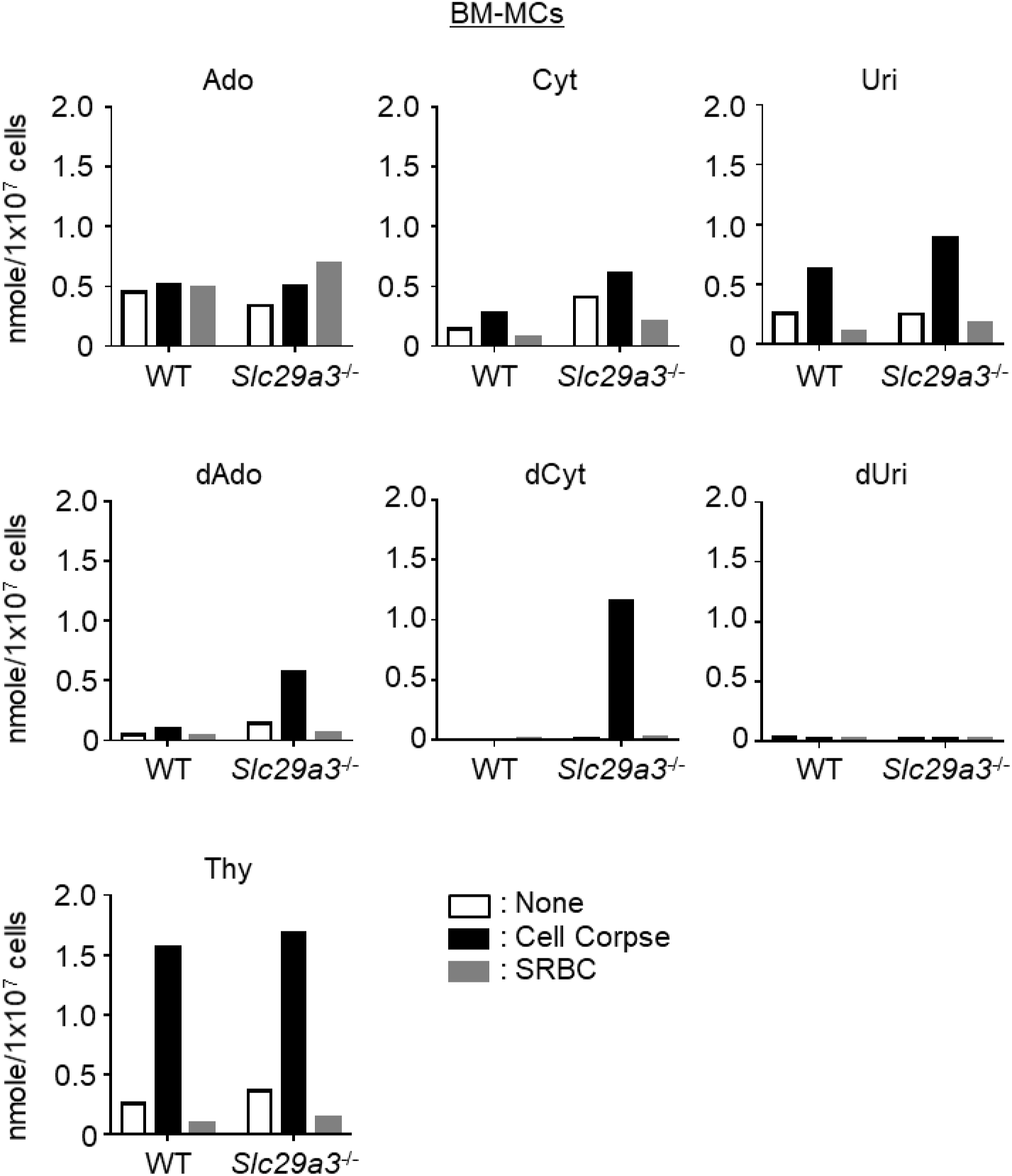
Nucleoside storage in macrophages after engulfment of cell corpse or SRBC. The amounts (nanomole) of accumulated nucleosides in 10^7^ cells of WT and *Slc29a3*^−/−^ BM-derived macrophages after 1d treatment with 10^8^ dying thymocytes (cell corpse) or 10^9^ sheep red blood cell (SRBC) are analyzed by LC-MS. The data for G and dG are shown in Fig. 2C.

**Figure S10.**
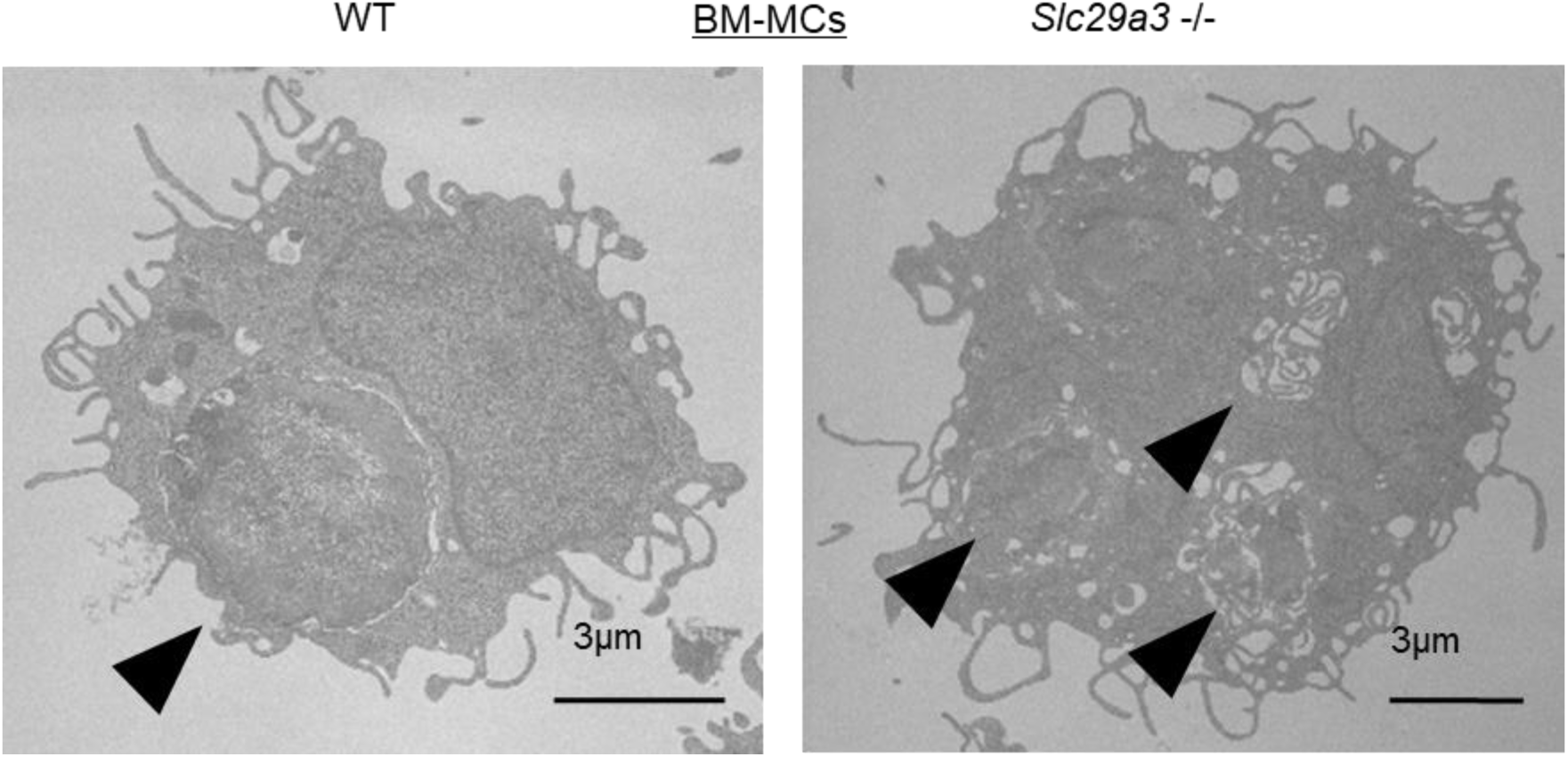
Unaltered phagosome formation in *Slc29a3*^−/−^ BM-macrophages. Electron Microscope images of WT and *Slc29a3*^−/−^ BM-macrophages (BM-MCs) having engulfed dying thymocytes. Arrow head, phagosomes.

**Figure S11.**
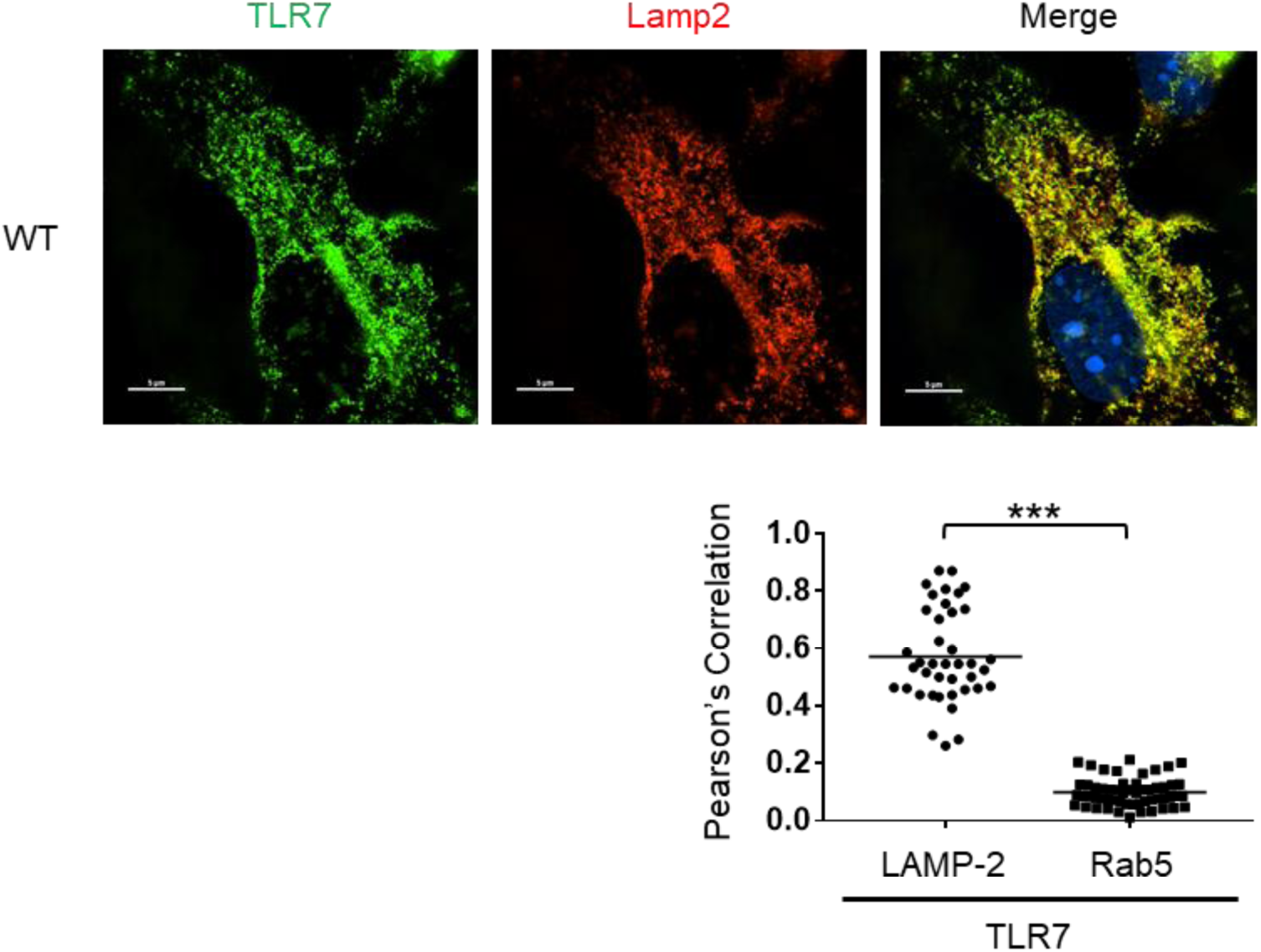
TLR7 is localized to LAMP-2-positive vesicles. WT BM-MCs were stained by antibodies to TLR7, an early endosome marker Rab5, and a lysosome marker LAMP-2. Statistical analyses were conducted to see colocalization between TLR7 and Rab5 or between TLR7 and LAMP-2. \*\*\**p*<0.001

**Figure S12.**
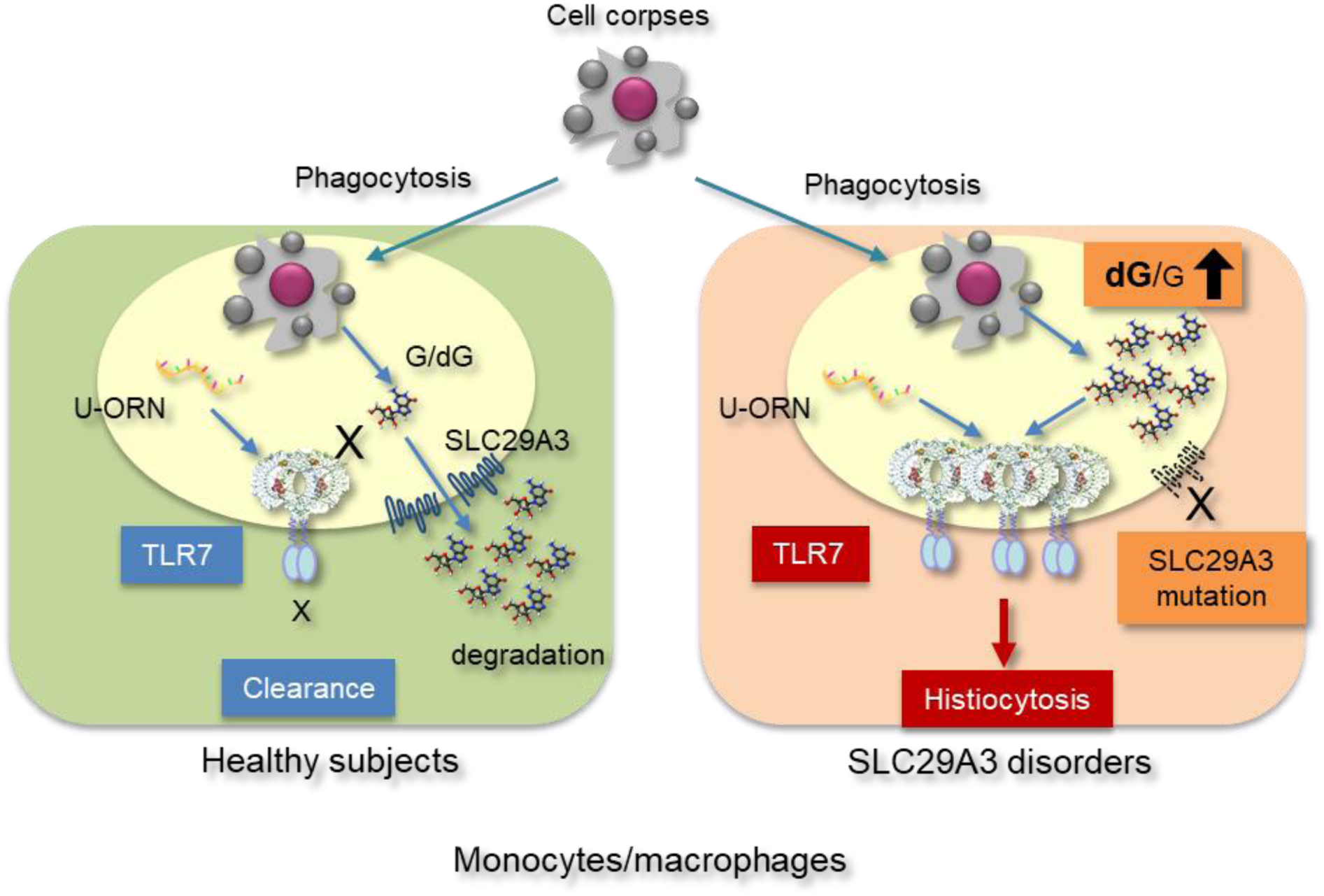
SLC29A3 prevents TLR7-dependent histiocytosis. Schematic representation of the mechanism silencing TLR7 responses to cell corpses by SLC29A3 in monocytes/macrophages. SLC29A3 is localized to phagosomes containing cell corpses and silences phagosomal TLR7 activation by tranporting G and dG out of phagosomes. TLR7 is recruited but not activated. In SLC29A3 disorders, dG and G accumulate and act on TLR7, leading to further TLR7 recruitment to phagosomes and TLR7 activation. TLR7 activation in phagosomes drives histiocytosis.

